# GLOF: A large-scale expert-curated benchmark dataset of gain-of-function and loss-of-function missense variants

**DOI:** 10.64898/2026.06.05.729843

**Authors:** Victor Maricato, David Schlesinger, Pedro Nuno de Souza Moura

## Abstract

Distinguishing loss-of-function (LOF) from gain-of-function (GOF) effects of missense variants is fundamental to understanding disease mechanisms and guiding therapeutic strategy, yet no large-scale, expert-curated benchmark has been publicly available for this task. Here we present GLOF (Gain and Loss Of Function), a dataset of 112,399 missense variants across 2,809 human genes, each classified as LOF, GOF, or neutral by board-certified clinical geneticists following ACMG guidelines. Pathogenic variants were sourced from ClinVar and annotated with their functional mechanism based on published functional studies, phenotype correlations, and established gene-disease relationships. Neutral variants were drawn from gnomAD v3.1 and validated against v4.1 using stringent population frequency filters. The dataset spans diverse protein families, includes 97 genes with bidirectional mechanisms (containing both LOF and GOF variants), and has been validated against well-characterized variants in the literature. GLOF is publicly available on Kaggle (https://www.kaggle.com/datasets/maricatovictor/loss-and-gain-of-function-variants) and Hugging Face (https://huggingface.co/datasets/victormaricato/glof), and provides a standardized resource for developing and benchmarking computational methods that predict variant functional mechanisms.

## 1 Background & Summary

The clinical interpretation of missense variants remains a central bottleneck in precision medicine [1, 2]. Current computational tools can predict whether a variant is pathogenic [3–7], but they largely fail to predict *how* it causes disease: whether through a loss-of-function (LOF) or gain-of-function (GOF) mechanism. This distinction has direct therapeutic consequences. LOF variants may require protein replacement or gene augmentation therapies, whereas GOF variants typically call for targeted inhibitors or allele-specific silencing [8, 9].

Several genes illustrate this point by producing opposite phenotypes depending on variant type. In KCNJ11, which encodes the pore-forming subunit (Kir6.2) of the ATP-sensitive potassium channel in pancreatic *β*-cells, GOF variants such as R201C reduce ATP sensitivity and hold the channel open, causing neonatal diabetes, whereas LOF variants such as V290M lock the channel closed, causing congenital hyperinsulinism [10, 11] (Fig. 1). In FGFR3, GOF variants cause achondroplasia (short stature) through constitutive kinase activation, while the rare LOF variant T546K causes CAT-SHL syndrome, characterized by tall stature [12, 29]. These cases show that predicting the direction of functional change, not merely its presence, is essential for precision therapeutics.

**Fig. 1.**
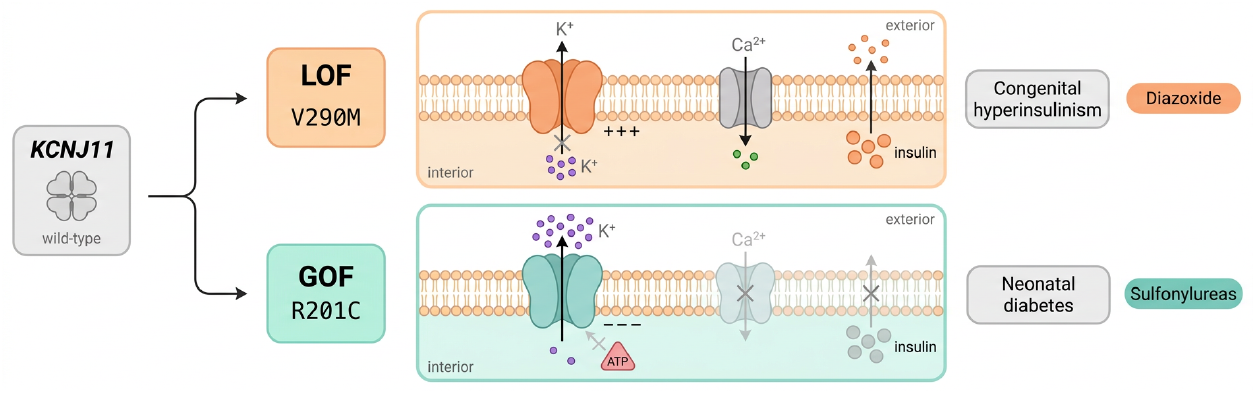
Bidirectional disease mechanisms in KCNJ11. The ATP-sensitive K^+^ (K_ATP_) channel in pancreatic *β*-cells illustrates how opposing missense variants in the same gene produce opposite clinical phenotypes. LOF variant V290M locks the channel closed, preventing K^+^ efflux and causing persistent membrane depolarization, excessive insulin secretion, and congenital hyperinsulinism (treated with diazoxide). GOF variant R201C keeps the channel constitutively open, maintaining membrane hyperpolarization, blocking insulin secretion, and causing neonatal diabetes (treated with sulfonylureas)[10, 11]. Purple circles represent K^+^ ions; green, Ca^2+^; peach, insulin granules.

Despite this need, computational methods for LOF/GOF prediction have been held back by the lack of large-scale, curated benchmark data. Existing resources focus on pathogenicity prediction without mechanism classification [13], provide gene-level rather than variant-level annotations [14], or cover only small numbers of well-studied proteins [15]. Gerasimavicius et al. [16] curated a dataset of 3,685 variants for structural analysis of LOF, GOF, and dominant-negative effects, but this remains too small for training machine learning models. The recent LoGoFunc method [15] was trained on datasets derived from automated database mining, which may propagate annotation errors and biases from the source databases.

Here we present GLOF (Gain and Loss Of Function), a benchmark dataset of 112,399 missense variants across 2,809 human genes, each classified as LOF, GOF, or neutral by board-certified clinical geneticists. The dataset was curated at Mendelics Análise Genômica^1^, one of Latin America’s largest clinical genomics laboratories, which has sequenced more than one million individuals. The annotation process integrated ClinVar pathogenicity classifications [13], published functional studies (e.g., [10, 11]), established gene-disease relationships (e.g., [12]), and expert clinical judgment following American College of Medical Genetics and Genomics (ACMG) and Association for Molecular Pathology (AMP) guidelines [1]. Variants with conflicting evidence were excluded to ensure label quality.

GLOF is, to the best of our knowledge, the largest publicly available expert-curated dataset of missense variants with functional mechanism labels. It provides a standardized benchmark for developing and evaluating computational tools that predict LOF versus GOF effects, a capability that grows more important as genomic sequencing becomes part of routine clinical care.

## 2 Methods

### 2.1 Data sources

The GLOF dataset integrates variants from two primary public databases, supplemented by the internal clinical database of Mendelics Análise Genômica (Fig. 2).

**Fig. 2.**
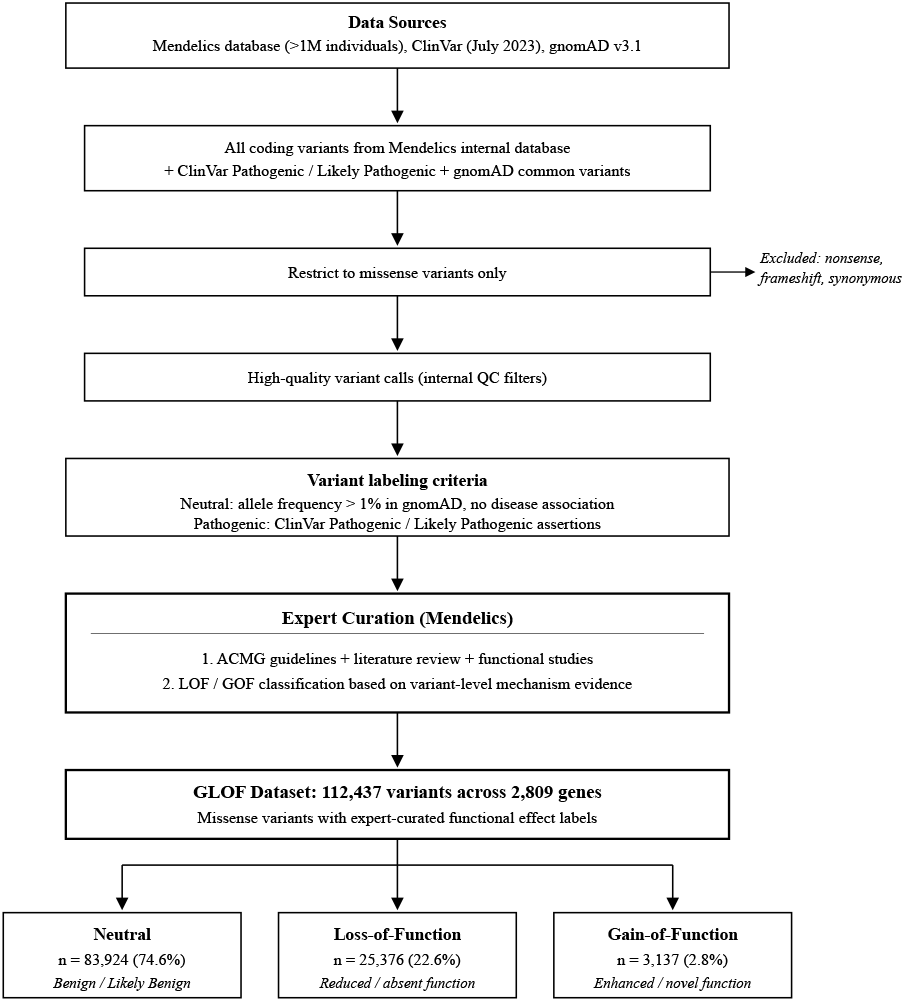
Dataset curation workflow. Variants were aggregated from the Mendelics internal database (*>*1 million individuals) and ClinVar, filtered for quality and variant type (missense only), and subjected to expert curation by board-certified clinical geneticists. Functional classifications were assigned following ACMG guidelines based on literature review of functional studies and phenotype correlations: Gain-of-Function (GOF) for variants with evidence of increased or novel protein activity, Loss-of-Function (LOF) for variants with evidence of reduced or absent activity, including haploin-sufficiency, and Neutral for benign variants identified through population frequency filters.

**Fig. 3.**
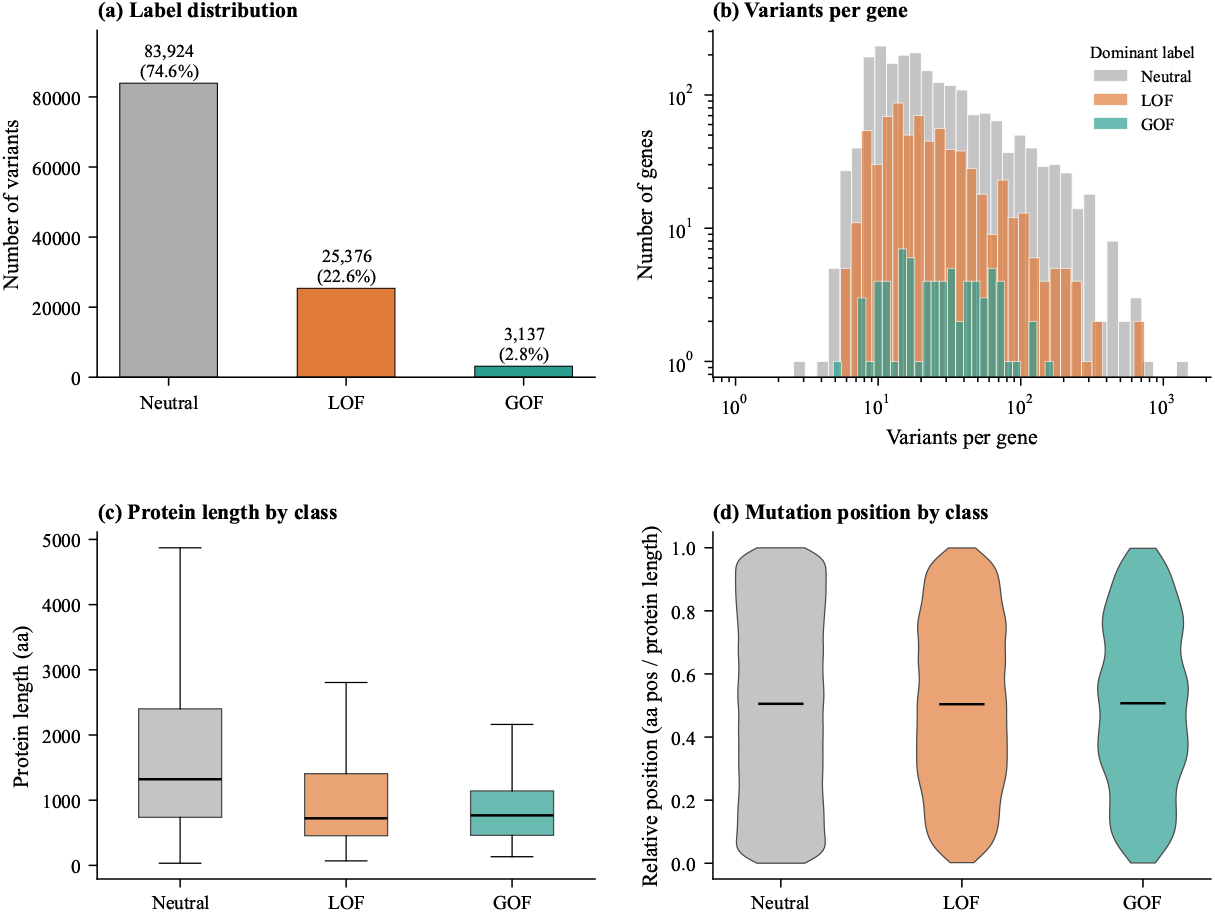
Overview of the GLOF dataset. (**a**) Distribution of variant classes: neutral (83,902; 74.6%), LOF (25,368; 22.6%), and GOF (3,129; 2.8%). (**b**) Distribution of variants per gene (log scale), colored by the dominant label in each gene. (**c**) Protein length distribution across variant classes, showing comparable length distributions. (**d**) Relative mutation position (normalized to protein length) for each class, showing uniform distribution across all three classes.

**Pathogenic variants** were sourced from ClinVar (July 2023 release) [13] and cross-referenced against the March 2026 release; variants with reclassified or conflicting evidence were excluded. Only variants classified as “Pathogenic” or “Likely Pathogenic”^2^ with at least one supporting submission were retained. Only missense variants (single amino acid substitutions) in protein-coding genes were retained. The Mendelics internal database, comprising sequencing data from over one million individuals, was used to cross-reference variant calls and confirm quality.

**Neutral variants** were drawn from the Genome Aggregation Database (gnomAD) v3.1 [18] and validated against v4.1 allele frequencies, selecting missense variants with allele frequency (AF) greater than 1% in at least one population and no reported disease association. This stringent frequency threshold ensures that neutral variants are common, well-tolerated polymorphisms rather than rare variants of uncertain significance.

### 2.2 Expert curation process

The functional annotation of variants was performed by clinical geneticists at Mendelics Análise Genômica, targeting genes with well-characterized functional mechanisms in the literature. For each pathogenic variant, the curation process involved:

1. **Gene-level mechanism assessment**: The established disease mechanism for each gene was determined from the literature. For genes with unidirectional mechanisms (e.g., BRCA1 exclusively causes disease through LOF; BRAF exclusively through GOF), this gene-level knowledge informed the variant labels.
2. **Variant-level review**: Each variant was analyzed by a single specialist who assessed whether the variant’s functional effect was consistent with the gene’s known mechanism. For genes with bidirectional mechanisms (e.g., KCNJ11, where different variants cause either neonatal diabetes via GOF or hyperinsulinism via LOF), variants were individually classified based on published functional studies and phenotype correlations. Variants with uncertain or insufficient evidence for mechanism classification were excluded from the dataset.

The classification criteria were:

- **Loss-of-function (LOF)**: Evidence supports reduced or absent protein function, including haploinsufficiency, null alleles, and dominant-negative effects that ultimately reduce pathway activity.
- **Gain-of-function (GOF)**: Evidence supports enhanced protein activity, constitutive activation, novel function, or altered specificity.
- **Neutral**: Population frequency data (gnomAD AF *>* 1%) or functional evidence indicates no pathogenic effect. Corresponds to ClinVar “Benign” or “Likely Benign” classifications.

The curation followed the ACMG/Association for Molecular Pathology (AMP) standards and guidelines for sequence variant interpretation [1], integrating multiple evidence types including population frequency data, functional studies, computational predictions, segregation data, and phenotype correlations.

### 2.3 Handling of dominant-negative variants

Dominant-negative (DN) variants, which interfere with the function of the wild-type protein product (e.g., through oligomeric poisoning), were classified as LOF in the GLOF dataset. This decision reflects the fact that DN variants ultimately result in reduced pathway activity, even though their molecular mechanism differs from simple haploinsufficiency [16, 19].

This classification is consistent with clinical practice, where DN and LOF variants in the same gene typically produce overlapping phenotypes [16, 17]. For example, in FBN1 (Marfan syndrome), approximately 59% of pathogenic variants act through a DN mechanism and 41% through haploinsufficiency [20], yet both produce the same clinical syndrome and are managed similarly. The GLOF dataset contains 688 LOF variants in FBN1, encompassing both DN and haploinsufficiency mechanisms.

We note this as a deliberate design choice rather than a limitation. Users requiring DN as a separate category may cross-reference GLOF with the dataset of Gerasimavicius et al. [16], which provides DN annotations for a subset of variants. Recent work has shown that DN variants have structurally distinct properties: they are enriched at protein-protein interfaces and exhibit milder structural perturbation than LOF variants [16, 17]. Future versions of GLOF could benefit from incorporating DN as a fourth category.

### 2.4 Quality control

Several quality control measures were applied during curation:

- **Variant call quality**: Only high-confidence variant calls were included, using quality metrics from both the Mendelics pipeline and ClinVar submission data.
- **Missense filtering**: Only single-nucleotide variants resulting in amino acid substitutions were retained; frameshifts, nonsense, synonymous, and splice variants were excluded.
- **Conflict resolution**: Variants with uncertain or conflicting mechanistic evidence were excluded, prioritizing precision over recall.
- **Gene-level validation**: Genes were required to have established functional characterization in the literature. Genes with insufficient functional evidence were excluded.

## 3 Data Records

The GLOF dataset is publicly available on Kaggle (https://www.kaggle.com/datasets/maricatovictor/loss-and-gain-of-function-variants) and Hugging Face (https://huggingface.co/datasets/victormaricato/glof). The dataset is provided as a tab-separated file with the following fields:

### 3.1 Dataset summary statistics

The dataset contains 112,399 missense variants across 2,809 genes (Table 2). The class distribution reflects the biological reality that most observed variants are benign and that GOF mechanisms are rarer than LOF mechanisms in human disease [17].

**Table 1.** GLOF dataset schema. Description of each field in the dataset.

| Field | Type | Description |
| --- | --- | --- |
| VARIANTKEY | String | Unique variant identifier in the format `chr-position-ref-alt` (GRCh38 coordinates) |
| LABEL | String | Functional classification: “Neutral”, “LOF”, or “GOF” |
| ENSG | String | Ensembl gene identifier |
| GENE_SYMBOL | String | HUGO Gene Nomenclature Committee (HGNC) gene symbol |
| AA_POSITION | Integer | Position of the amino acid substitution in the canonical protein sequence |
| PROTEIN_REF | Character | Reference (wild-type) amino acid at the substitution position |
| PROTEIN_ALT | Character | Alternate (mutant) amino acid at the substitution position |

**Table 2.**
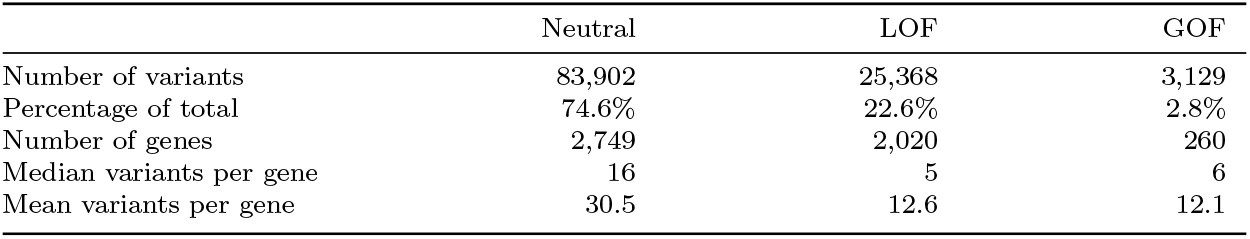
Summary statistics of the GLOF dataset.

### 3.2 Gene-level label patterns

Gene-level label patterns fall into four categories that reflect the underlying biology of disease mechanisms (Fig. 4a):

**Fig. 4.**
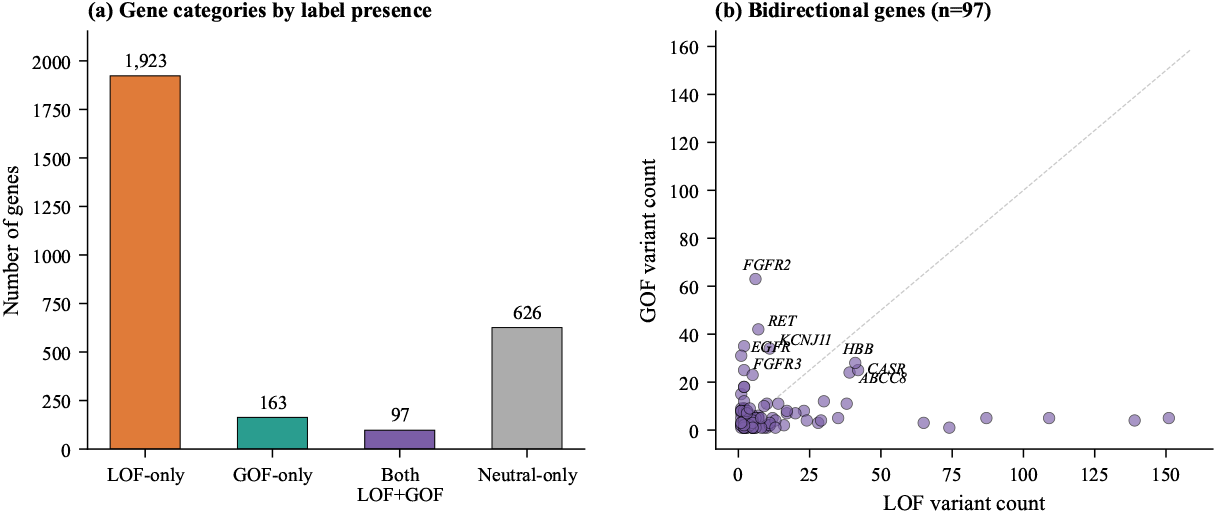
Gene-level label patterns. (**a**) Distribution of genes by label presence: 1,923 genes have only LOF pathogenic variants, 163 have only GOF, 97 have both (bidirectional), and 626 have only neutral variants. (**b**) For the 97 bidirectional genes, scatter plot of LOF versus GOF variant counts. Key genes with well-characterized bidirectional mechanisms are labeled. The dashed line indicates equal LOF and GOF counts.

- **LOF-only genes** (1,923 genes): Genes where all pathogenic variants act through LOF. These include well-known tumor suppressors (BRCA1, TP53) and structural proteins (FBN1, COL3A1).
- **GOF-only genes** (163 genes): Genes where all pathogenic variants act through GOF. These include oncogenes (BRAF, KRAS) and constitutively activating ion channel variants.
- **Bidirectional genes** (97 genes): Genes containing both LOF and GOF variants, where different mutations produce opposite functional effects. These represent the most challenging cases for computational prediction.
- **Neutral-only genes** (626 genes): Genes with only neutral variants in the dataset, reflecting genes where no pathogenic missense variants were available in the source databases.

The 97 bidirectional genes are of particular biological interest, as they demonstrate that the functional effect of a missense variant cannot be inferred solely from the gene in which it occurs. Prominent examples include:

- **KCNJ11** (34 GOF, 11 LOF): GOF variants reduce ATP sensitivity, causing neonatal diabetes; LOF variants inactivate the channel causing congenital hyperinsulinism [10, 11].
- **CASR** (25 GOF, 42 LOF): GOF variants increase calcium sensitivity, causing autosomal dominant hypocalcemia; LOF variants decrease sensitivity causing familial hypocalciuric hypercalcemia [21].
- **HBB** (28 GOF, 41 LOF): GOF variants such as E7V (sickle hemoglobin) confer neomorphic polymerization of deoxy-hemoglobin; LOF variants cause beta-thalassemia through reduced or absent functional beta-globin [33, 34].
- **RET** (42 GOF, 7 LOF): GOF variants cause multiple endocrine neoplasia type 2 (MEN2) through constitutive kinase activation; LOF variants cause Hirschsprung disease through impaired neural crest migration [22, 23].
- **FGFR3** (48 GOF, 2 LOF): GOF variants such as G380R cause achondroplasia; the rare T546K LOF variant causes the opposite phenotype of tall stature (CATSHL syndrome) [12, 27].

Figure 5 illustrates the spatial distribution of LOF and GOF variants on the AlphaFold-predicted structures [36, 37] of three representative bidirectional genes. In (**A**), GOF variants concentrate in the cytoplasmic N-terminal and C-terminal domains involved in ATP-binding and channel gating (23 of 26 GOF variants), while LOF variants are more evenly distributed across all domains. In (**B**), mutations span the large extracellular Venus flytrap domain and 7-TM region. In (**C**), the compact globin fold shows extensive interleaving of LOF and GOF sites, consistent with the distinct molecular mechanisms underlying beta-thalassemia (LOF) and sickle cell disease (GOF).

**Fig. 5.**
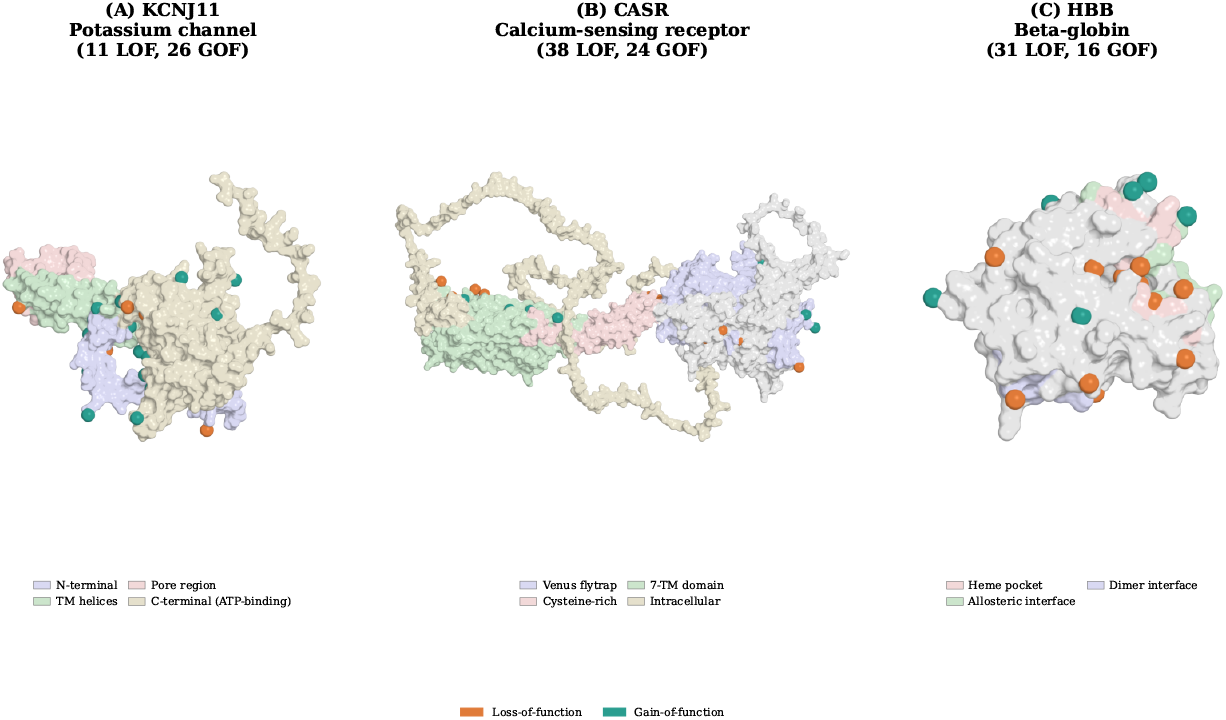
Spatial distribution of LOF and GOF variants in bidirectional genes. AlphaFold-predicted structures [36, 37] of (**A**) KCNJ11 (potassium channel, UniProt Q14654), (**B**) CASR (calcium-sensing receptor, UniProt P41180), and (**C**) HBB (beta-globin, UniProt P68871) shown as molecular surfaces, with LOF variant positions marked in orange and GOF variant positions in teal. Surface regions are colored by functional domain as annotated in UniProt [38] (see per-protein legends). In (**A**), GOF variants concentrate in the cytoplasmic N-terminal and C-terminal (ATP-binding) domains, while LOF variants are distributed across all domains including the pore region; the R201C (GOF) and V290M (LOF) variants illustrated in Fig. 1 both map to the C-terminal domain. In (**B**), both LOF and GOF variants span the Venus flytrap and 7-TM domains. In (**C**), the compact globin fold shows interleaving of LOF and GOF sites near the heme pocket and subunit interfaces.

### 3.3 Amino acid substitution patterns

Pathogenic variants (both LOF and GOF) are enriched for class-changing substitutions relative to neutral variants. Among LOF variants, 74.8% involve a change in amino acid physicochemical class (e.g., polar to nonpolar), compared to 72.1% for GOF and 60.7% for neutral (*p <* 10^*−16*^, *χ*^2^ test; Fig. 6c). This is consistent with the expectation that altering protein function typically requires substantial changes to local physicochemical properties.

**Fig. 6.**
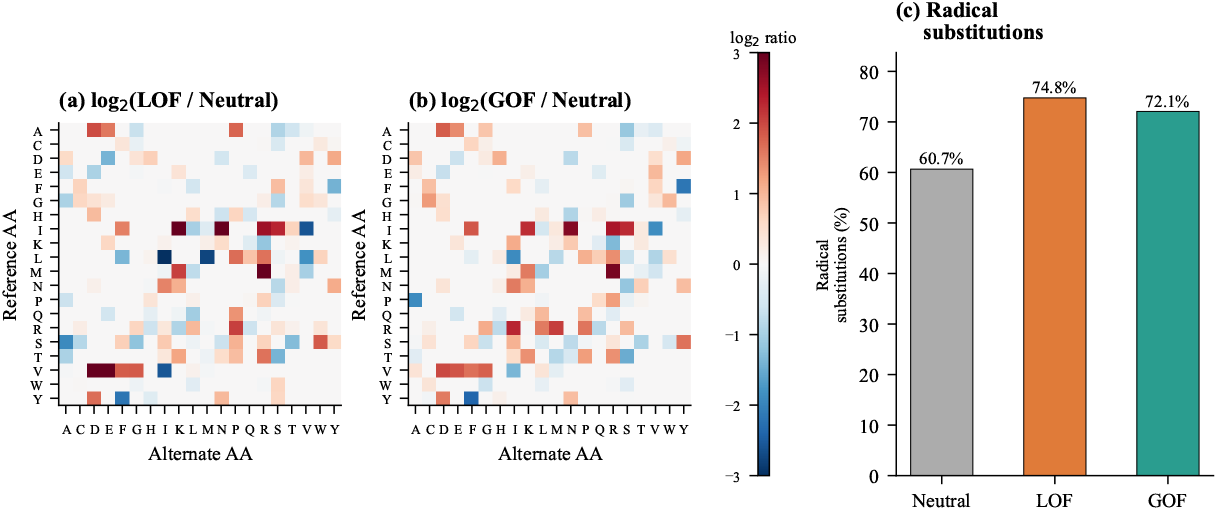
Amino acid substitution patterns. (**a**) Log_2_ ratio of LOF to neutral substitution frequencies across all 20 *×*20 amino acid pairs. Red indicates enrichment in LOF; blue indicates depletion. (**b**) Same analysis for GOF versus neutral. (**c**) Percentage of radical substitutions by variant class. Both LOF (74.8%) and GOF (72.1%) are significantly enriched for radical substitutions compared to neutral variants (60.7%; *p <* 10^*−16*^).

**Fig. 7.**
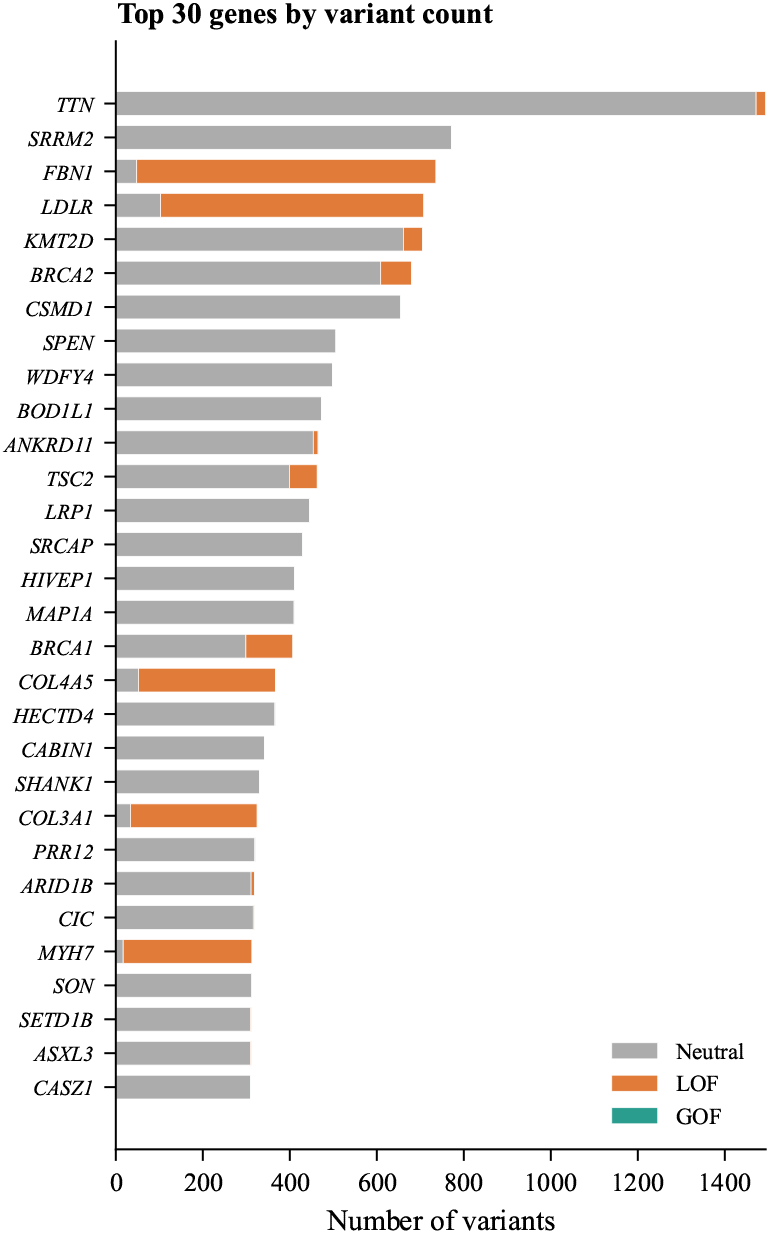
Top 30 genes by total variant count. Horizontal stacked bars show the contribution of each label class. Clinically important genes such as TTN (titin), FBN1 (fibrillin-1), LDLR (LDL receptor), BRCA1/2, and structural collagens (COL4A5, COL3A1) are well-represented.

## 4 Technical Validation

### 4.1 Cross-validation with literature

To assess label accuracy, we cross-referenced GLOF annotations against well-characterized variants with published functional evidence. Table 3 presents representative examples from bidirectional genes, where the same gene contains both LOF and GOF variants with different, experimentally validated mechanisms.

**Table 3.** Validation of GLOF labels against published functional evidence. All 13 cross-checked variants from bidirectional genes matched published literature.

| Gene | Variant | GLOF | Known mechanism | Reference |
| --- | --- | --- | --- | --- |
| FGFR3 | G380R | GOF | Constitutive kinase activation (achondroplasia) | [27] |
| FGFR3 | K650E | GOF | ~100× kinase activation | [28, 29] |
| FGFR3 | T546K | LOF | CATSHL syndrome (tall stature) | [12] |
| KCNJ11 | V59M | GOF | Reduced ATP sensitivity (DEND) | [10] |
| KCNJ11 | R201C | GOF | Neonatal diabetes | [10] |
| KCNJ11 | V290M | LOF | Channel inactivation (hyperinsulinism) | [11] |
| RET | M918T | GOF | Constitutive kinase activation (MEN2B) | [23] |
| CASR | A843E | GOF | Increased Ca<sup>2+</sup> sensitivity | [21] |
| CASR | R185Q | LOF | Decreased Ca<sup>2+</sup> sensitivity (NSHPT) | [21] |
| HBB | E7V | GOF | Neomorphic polymerization (sickle cell) | [24] |
| ABCC8 | R1182Q | GOF | Enhanced channel opening (neonatal diabetes) | [25] |
| ABCC8 | R526C | LOF | Reduced channel activity (hyperinsulinism) | [26] |

All 13 variants examined matched their published functional characterization, supporting the accuracy of the expert curation process. This validation set includes variants from genes where different substitutions at the same protein, or even the same amino acid position, have opposite effects, confirming that GLOF captures genuine variant-level functional distinctions.

### 4.2 Same-position, opposite-effect variants

The strongest evidence for variant-level (rather than gene-level) curation comes from cases where different amino acid substitutions at the same genomic position received opposite labels:

- **CASR position 127**: E127A is labeled GOF; E127G is labeled LOF (same genomic coordinate chr3:122257275).
- **CASR position 221**: P221L is labeled GOF; P221Q is labeled LOF (same genomic coordinate chr3:122261697).
- **HBB position 24**: V24F is labeled GOF; V24I is labeled LOF (same genomic coordinate chr11:5226952).

These cases confirm that the curation resolved functional effects at the individual variant level, even when two substitutions at the same position produce opposite outcomes owing to the distinct physicochemical properties of the substituted amino acids.

### 4.3 Comparison with existing resources

Table 4 compares GLOF with other resources that include LOF/GOF annotations. GLOF provides substantially more variants than previous manually curated datasets while maintaining expert-level annotation quality.

**Table 4.** Comparison of GLOF with existing LOF/GOF variant resources.

| Resource | Variants | Genes | Curation | LOF/GOF | Neutral |
| --- | --- | --- | --- | --- | --- |
| GLOF (this work) | 112,399 | 2,809 | Expert | Yes | Yes |
| Gerasimavicius 2022 [16] | 3,685 | — | Semi-auto | Yes+DN | No |
| LoGoFunc training [15] | ~20,000 | — | Automated | Yes | Yes |
| ClinVar [13] | >2M | >40,000 | Submitter | No<sup>1</sup> | Yes |

<sup>1</sup>ClinVar provides pathogenicity classification but does not distinguish LOF from GOF mechanism.

### 4.4 Label distribution analysis

The class imbalance in GLOF (74.6% neutral, 22.6% LOF, 2.8% GOF) reflects known properties of the human variant landscape. Most observed variants are benign [18], and among pathogenic variants, LOF mechanisms are roughly eight times more prevalent than GOF mechanisms. This ratio is consistent with Gerasimavicius et al. [17], who found that LOF accounts for the majority of disease mechanisms in Mendelian disorders, with GOF and dominant-negative mechanisms being considerably rarer.

## 5 Usage Notes

### 5.1 Limitations and considerations

Users should be aware of the following considerations when using GLOF:

1. **Dominant-negative variants**: DN variants are classified as LOF (Section 2.3). The LOF class therefore contains a mixture of haploinsufficiency, dominant-negative, and recessive LOF variants, which may have distinct structural and evolutionary signatures [16].
2. **Gene coverage bias**: The dataset focuses on genes with well-characterized mechanisms, which biases toward clinically studied genes. Genes associated with rare diseases or those with limited functional characterization may be underrepresented.
3. **Ascertainment bias**: Pathogenic variants are ascertained from clinical databases (ClinVar), which are enriched for variants observed in diagnostic settings. This may underrepresent pathogenic variants in populations not well served by genetic testing.
4. **Neutral variant definition**: Neutral variants are defined by population frequency (*>* 1% in gnomAD), not by functional assay. While high-frequency variants are overwhelmingly benign, a small number may have mild functional effects that, although compatible with normal health, are not truly functionally neutral [18].
5. **Binary LOF/GOF classification**: The LOF/GOF distinction simplifies a continuous spectrum of variant effects. Some variants may have partial LOF, context-dependent effects, or mixed LOF/GOF properties that are not captured in the current classification.

## Acknowledgements

We thank the clinical genetics team at Mendelics Análise Genômica, in particular Prof. Dr. Fernando Kok, for their work in curating the variant annotations. We also thank the maintainers of ClinVar and gnomAD for making their data publicly available.

## Declarations

### Funding

Dataset curation was supported by Mendelics Análise Genômica.

### Competing interests

D.S. is CEO of Mendelics Análise Genômica. Other authors declare no competing interests.

### Ethics approval

Not applicable. This study uses previously collected, de-identified genetic variant data from public databases (ClinVar, gnomAD). No individual-level patient data was used.

### Consent for publication

Not applicable.

### Data availability

The GLOF dataset is publicly available on Kaggle at https://www.kaggle.com/datasets/maricatovictor/loss-and-gain-of-function-variants and on Hugging Face at https://huggingface.co/datasets/victormaricato/glof.

### Author contributions

D.S. led the clinical curation of the dataset. V.M. performed the computational analysis, validation, and wrote the manuscript. P.N.M. supervised the computational methodology. All authors reviewed and approved the final manuscript.

## Footnotes

1 https://www.mendelics.com/

2 ClinVar clinical significance classifications: https://www.ncbi.nlm.nih.gov/clinvar/docs/clinsig/

## Notes

### Competing Interest Statement

David Schlesinger (D.S.) is CEO of Mendelics Analise Genomica. The other authors declare no competing interests.

